# Identification of functional variants for platelet *CD36* expression by Massively Parallel Reporter Assay

**DOI:** 10.1101/550871

**Authors:** Namrata Madan, Andrew R. Ghazi, Xianguo Kong, Edward S. Chen, Chad A. Shaw, Leonard C. Edelstein

**Affiliations:** Cardeza Foundation for Hematologic Research/Department of Medicine, Sidney Kimmel Medical School, Thomas Jefferson University, Philadelphia, PA, USA; Department of Quantitative and Computational Biosciences, Baylor College of Medicine, Houston, TX, USA; Department of Molecular and Human Genetics, Baylor College of Medicine, Houston, TX, USA; Department of Statistics, Rice University, Houston, TX, USA

**Keywords:** CD36, eQTL, platelets, gene expression, genetic variation

## Abstract

CD36 is a platelet membrane glycoprotein whose engagement with oxidized low-density lipoprotein (oxLDL) results in platelet activation. The CD36 gene has been associated with platelet count, platelet volume, as well as lipid levels and CVD risk by genome-wide association studies. Platelet CD36 expression levels have been shown to be associated with both the platelet oxLDL response and an elevated risk of thrombo-embolism. Several genomic variants have been identified as associated with platelet CD36 levels, however none have been conclusively demonstrated to be causative. We screened 81 expression quantitative trait loci (eQTL) single nucleotide polymorphisms (SNPs) associated with platelet *CD36* expression by a Massively Parallel Reporter Assay (MPRA) and analyzed the results with a novel Bayesian statistical method. Ten eQTLs located in a 35kb region upstream of the *CD36* transcriptional start site demonstrated significant transcription shifts between their minor and major allele in the MPRA assay. Of these, rs2366739 and rs1194196, separated by only 20bp, were confirmed by luciferase assay to alter transcriptional regulation. In addition, electromobility shift assays demonstrated differential DNA:protein complex formation between the two alleles of this locus. Furthermore, deletion of the genomic locus by CRISPR/Cas9 in K562 cells results in upregulation of CD36 transcription. These data indicate that we have identified a variant that regulates expression of *CD36*, which in turn affects platelet function. To assess the clinical relevance of our findings we used the PhenoScanner tool, which aggregates large scale GWAS findings; the results reinforce the clinical relevance of our variants and the utility of the MPRA assay. The study demonstrates a generalizable paradigm for functional testing of genetic variants to inform mechanistic studies, support patient management and develop precision therapies.

**Author Summary:** Platelets are anucleate cells that are best known as regulators of vascular hemostasis and thrombosis but also play important roles in cancer, angiogenesis, and inflammation. CD36 is a platelet surface marker that can activate platelet in response to oxidized low density lipoprotein (oxLDL). CD36 has been associated with numerous cardiovascular traits in human including blood lipid levels, platelet count, and cardiovascular disease prevalence in human genetic studies. Human variability in platelet CD36 levels are associated with the platelet response to oxLDL. However, the genetic mechanisms responsible for the variability of CD36 levels are unknown. We examined 81 genetic variants associated with *CD36* levels for functionality using a high-throughput assay. Of the ten variants that were identified in that assay, one doublet, rs2366739 and rs1194196, were confirmed using additional molecular and cellular assays. Deletion of the genomic region containing rs2366739 and rs1194196 resulted in overexpression of *CD36* in a cell culture system. This finding indicates a control locus which can serve as a potential target in modulating CD36 expression and altering platelet function in cardiovascular disease.

## Introduction

Cardiovascular disease (CVD) remains the number one cause of death globally [1]. Myocardial infarctions (MI) are acute events in CVD which are frequently the proximal causes of death or severe disability which are the result of platelet-rich thrombi [2]. Genome wide association studies (GWASs) have identified numerous common genetic variants associated with the risk of CVD and platelet function parameters, but these variants are usually not causative due to the resolution of the genotyping platforms used and genetic linkage. One of the genes identified by GWAS as associated with platelet count, lipid levels. and CVD is the platelet oxidized LDL (oxLDL) receptor, CD36 [3-5].

CD36 is a transmembrane protein belonging to the class B scavenger receptor family expressed in platelets and variety of other cells [6-8]. It binds to many ligands such as oxidized phospholipids (oxPL) and oxidized low-density lipoprotein (oxLDL) long-chain fatty acids [9]. In platelets, CD36 interaction with oxLDL and thrombospondin-1 (TSP1) triggers MAP and Src family kinase dependent signaling events leading to platelet activation, [10, 11] which also lead to increase in P-selectin expression and αIIbβ3 activation [10]. Deletion of CD36 in mice fed a high fat diet results in attenuation of the pro-thrombotic state and platelet hyper-activity [10].

CD36 deficiencies have been identified which result in increased risk of cardiomyopathy, hyperlipidemia and insulin resistance [12-15]. In type I deficiency, monocytes and platelets lack CD36 expression, whereas in type II only platelets lack CD36 expression. CD36 deficiency is more frequent in black and Asian populations. Our platelet transcriptomic data also show that platelet *CD36* RNA levels are lower in the black population and in women [16, 17]. The molecular mechanisms behind *CD36* deficiency have been attributed to variants causing defects in protein maturation or frameshift, resulting in an absence of protein [14, 18].

Among subjects without CD36 deficiency, there is a wide range of platelet CD36 surface expression and the level of CD36 correlated with reactivity to oxLDL [19]. Many genetic variants have already been reported to be associated with platelet CD36 expression, however, these variants span a large linked genomic area and no functional analysis has been carried out [19, 20]. We have previously reported platelet expression Quantitative Trait Loci (eQTLs) that associate single nucleotide polymorphisms (SNPs) with platelet RNA levels, indicating genetic variability effecting gene expression [21]. *CD36* is one of the 612 platelet-expressed RNAs whose abundance has significant genotypic associations. 81 eQTL SNPs located within a +/-100kb window of the CD36 gene are associated with platelet CD36 mRNA levels at a significance of P<1×10^−6^, spanning a range of 118kb.

We hypothesized that a parallel screening method would be more efficient and cost-effective to identify causal variants instead of a one-at-a-time approach. We used a massively parallel reporter assay (MPRA) to screen the 81 platelet eQTLs associated with *CD36* mRNA. We developed new statistical methods for MPRA analysis, and we were able to identify rs2366739 and rs1194196 as functional variants that alter transcriptional regulation. We further tested these MPRA-functional variants, showing significant transcription shift between the reference and alternate alleles by luciferase assays, electromobility shift assays (EMSA) and using CRISPR/Cas edited stable cell lines. Finally, we used the Phenoscaner GWAS aggregation tool to reinforce the clinical relevance of our functional variants. Using these approaches, we have identified genetic variants that modulate platelet CD36 expression and have clinical associations.

## Results

### Landscape of CD36 Variants

We have previously published the results of a cis-eQTL analysis of platelet gene expression [21]. SNPs found to be associated with platelet *CD36* mRNA expression are indicated by diamonds on the Manhattan plot in Fig 1. Ghosh et al. have also looked for associations between *CD36* SNPs and CD36 protein expression [19]. SNPs identified in that report are indicated in Fig 1 by squares, and SNPs that were identified both by our eQTL study and Ghosh et al. are indicated by upside-down triangles. Several *CD36* SNPs have been identified in GWAS studies to associate with platelet count or volume. The GWAS-identified SNPS rs6961069, rs13236689, rs2177616, and rs11764390 are also platelet *CD36* eQTLs and are indicated by triangles in Fig 1 [4, 22, 23]. rs139761834 (Fig. 1, circle) was identified by GWAS but not by eQTL analysis [23]. The genomic region containing these variants encompasses the 5’ end of the *CD36* gene and upstream sequence and is highly linked as indicated by the linkage disequilibrium plot in the bottom panel of Fig 1. Given that most data on CD36 deficiency and expression has been obtained from Japanese subjects, D’ values were calculated using the 1000 Genomes JPT population. This strong linkage makes identification of the functional variant difficult, and therefore functional analysis requires experimental testing of individual SNPs. The large genomic span and the number of *CD36* phenotype-linked variants necessitated a highly parallel approach for functional testing.

**Fig 1.**
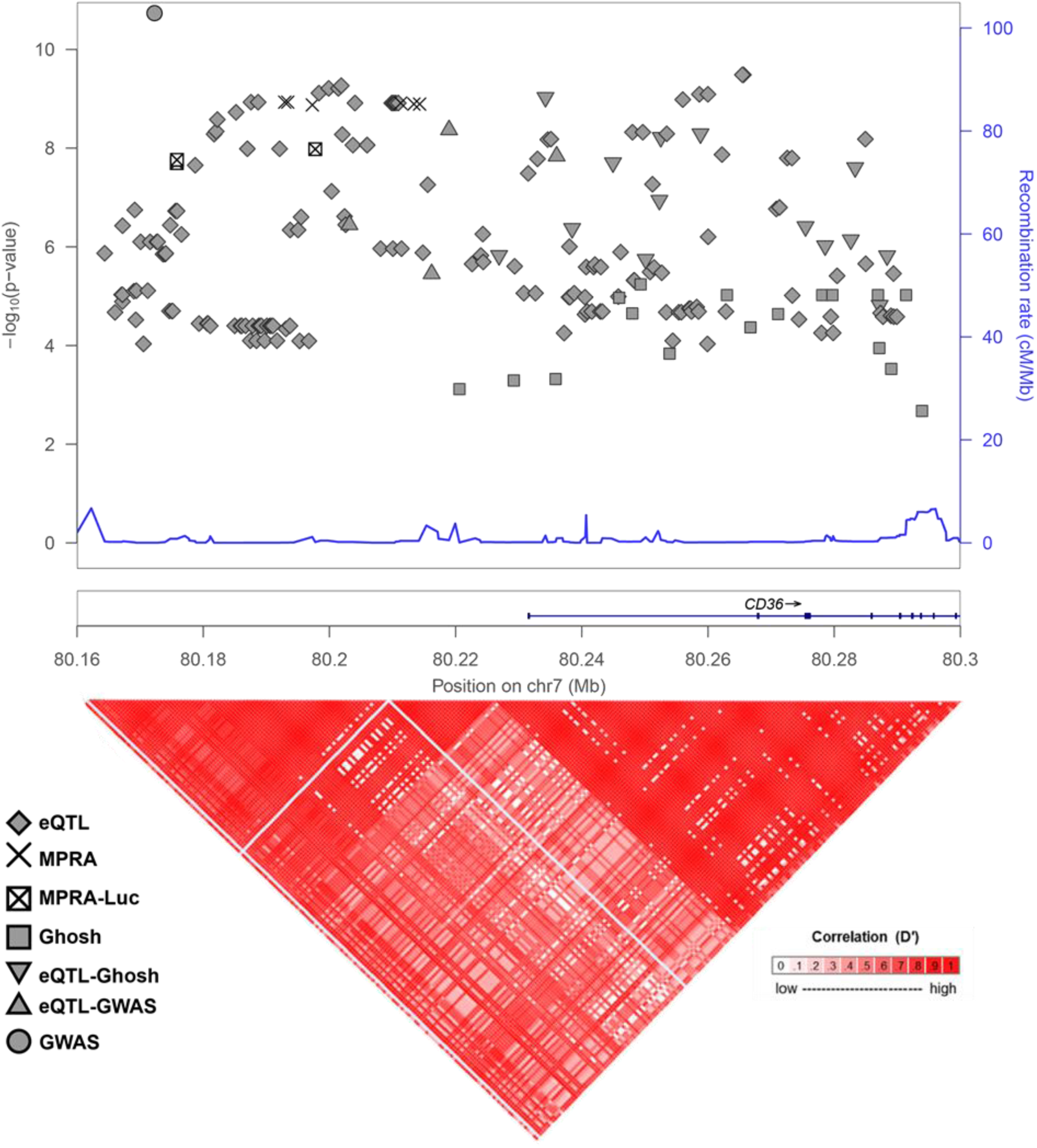
Genomic Context of *CD36* Variants. (Upper) Manhattan plot of *CD36* variants identified as platelet eQTLs only (diamonds), associated with CD36 surface levels (squares), GWAS-implicated (circle), eQTLs and CD36 surface level associated (upside down triangle), eQTLs and associated with GWAS-implicated (triangle), eQTLs and functional in MPRA assay (X’s), or eQTLs functional in MPRA assay and validated by luciferase (squared X). –log_10_ P-values are significance of eQTL association, if available, or from CD36 surface level association. GWAS-implicated variant (circle) is associated with platelet count (P=2 x 10^−17^) and vertical position does not represent significance. (Lower) LD-plot of genomic locus with D’ values calculated with 1000 genomes JPT (Japanese) population.

### Massively parallel reporter assay identifies CD36 SNPs with allelic difference in CD36 gene expression

We generated a library of plasmids in which a unique 10bp barcodes located downstream to a luciferase cassette were transcribed under the control of a 150bp genomic fragment containing a *CD36* platelet eQTL SNP. Each allele of each variant is associated with 40 unique barcodes in order to give high statistical power for detecting variant function despite variation in the NGS outputs. Transcription activity driven by each variant was then measured by quantifying RNA barcodes output after they have been “normalized” or jointly modeled against the plasmid DNA library barcode inputs.

We employed two statistical methods to analyze the result of the MPRA: a traditional method that involves computing variant activities defined by a ratio transformation of mRNA to DNA input and a second Bayesian method. The traditional method normalizes the counts for sample depth, removes barcodes with low representation in the plasmid library, and computes the activity as 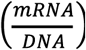 of each barcode in each allele in each transfection, and then uses a t-test and false discovery rate correction to compare the mean activity levels of alleles for each SNP. This approach yields 14 hits with Q <.05, including nine of the 10 controls (Table 1; Fig 2A) and five CD36 SNPs: rs2366739, rs940542, rs1093831, rs11464747, and rs6467258 (Table 1; Fig 2B). MPRA activities can occasionally defy the normality assumption underlying the t-test,[24] so we also employed the non-parametric Mann-Whitney U-test, under the same frequentist methodologic paradigm as the t-test approach. Analyzing the transcription activity level with a U-test revealed an additional four significant variants, rs1194196, rs6961069, rs819456, and rs819457 (Table 1; Fig 2C).

**Table 1.**
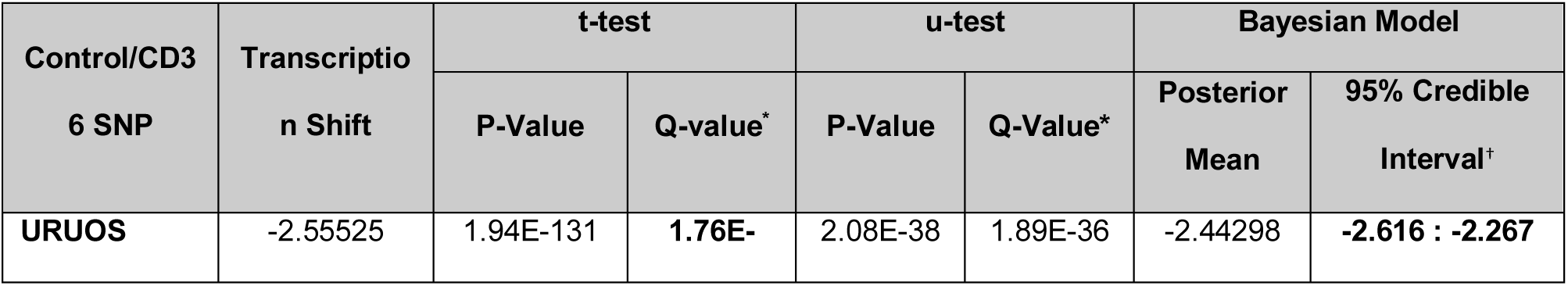
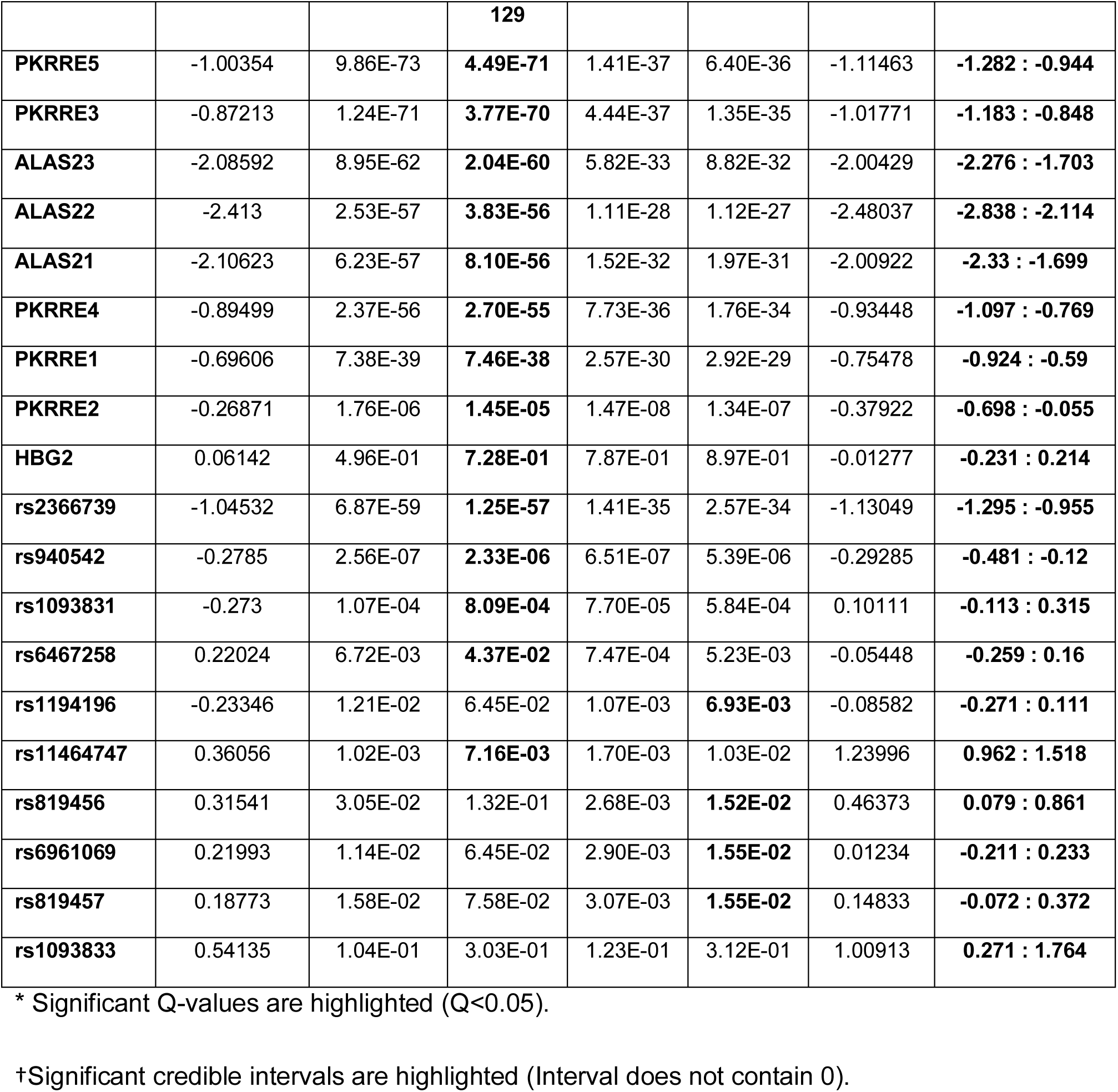
MPRA-functional *CD36* variants.

**Fig 2.**
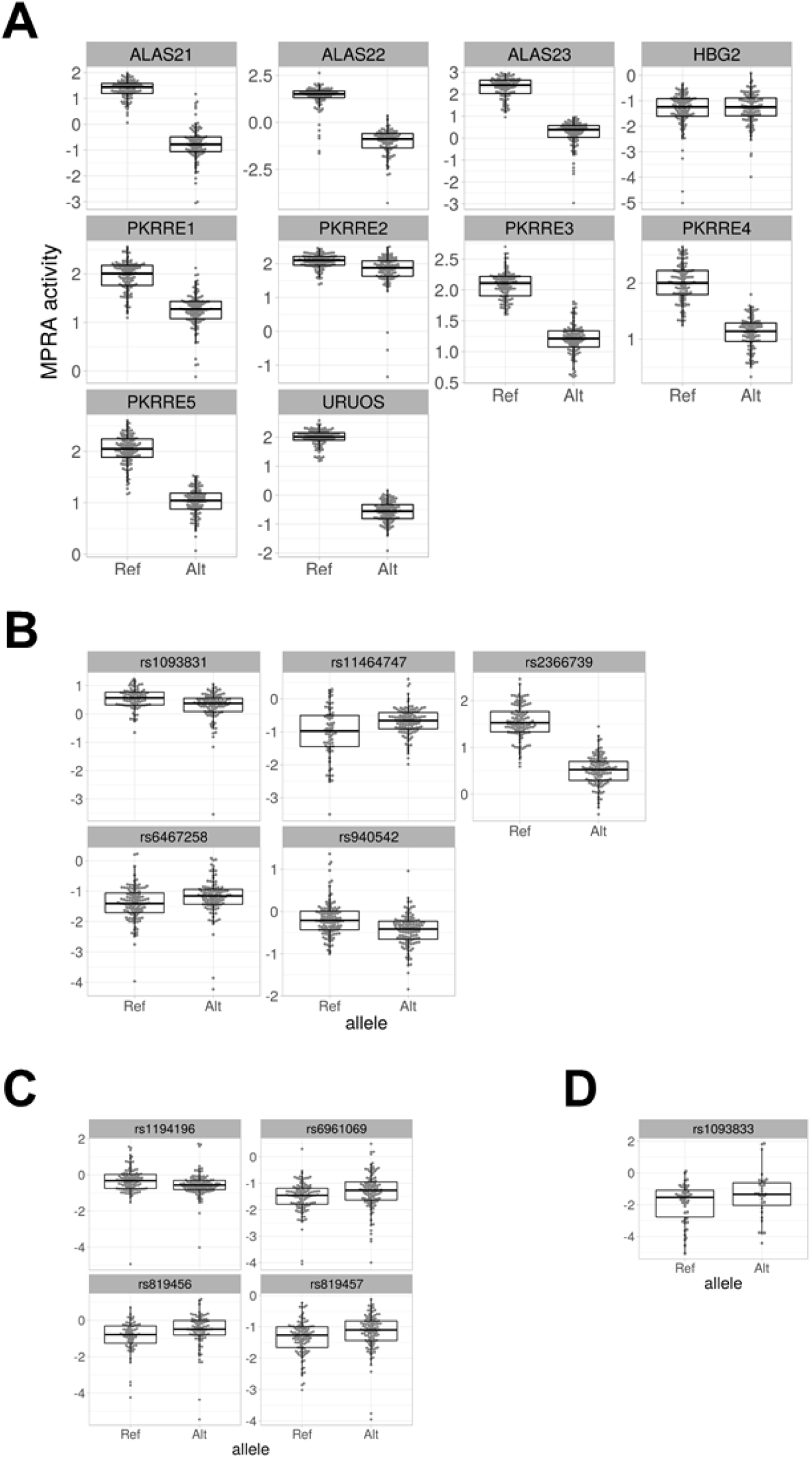
Results of MPRA assay of *CD36* eQTL SNPs. (A) MPRA activity of positive control constructs. (B) MPRA activity of *CD36* eQTL SNPs with significant transcription shifts identified by T-Test. (C) Additional *CD36* variants with significant transcription shifts identified by U-Test. (D) Additional *CD36* SNP with significant MPRA transcription shift identified by Bayesian analysis.

We sought to avoid two statistical limitations that lead to information and power loss in the MPRA experiment using the traditional analysis approach. First, transforming the data with a ratio removes the ability to model systematic effects of the DNA and RNA libraries. Second, discarding barcodes with low or 0 counts discards data that may be informative. Therefore, we also employed a Bayesian count model of the data generating process that models the NGS reads of each barcode observed from sequencing the plasmid library and from each transfection experiment as arising from coupled negative binomial distributions. The means of the negative binomial distributions are proportional to the depth of the sequencing of each sample and, in the case of RNA samples, the mean of the barcode’s DNA read count. Empirical gamma priors on the negative binomial parameters were estimated marginally across all SNPs in the assay. The log difference in the depth- and DNA-normalized RNA means gives a quantity comparable to the difference in mean activity (i.e. the ratios) analyzed under the t-test-based method. Thus, this model provides a posterior on transcription shift for each SNP after directly accounting for more sources of variation and more data from the MPRA experiment than the traditional approach. We identify a SNP in question as a functional hit if a 95% credible interval for the posterior distribution of the transcription shift excludes 0. This process yields 19 hits, including the same nine of the ten controls and ten CD36 SNPs, the nine listed above plus an additional variant, rs1093833 (Table 1; Figs 2D and S5) that was not identified by the frequentist approaches. As shown in Fig 1, (MPRA positive hits indicated by X’s) these SNPs are in high LD with one another, indicating close physical proximity in what is likely the regulatory region of the *CD36* gene. The complete analysis results of the tested MPRA variants and controls is given in Tables S4 and S5.

### Validation of differential enhancer activity of rs2366739-rs1193196 locus

To verify the transcription shifts identified by the MPRA, we tested three of the controls, the two most significant MPRA hits by t-test (rs2366739 and rs940542), the additional two most significant MPRA hits identified by U-test (rs1193196 and rs819456), and the additional significant MPRA hit identified by Bayesian analysis (rs1093833) by reporter assay. Reporter plasmids containing reference or alternate alleles of the eQTLs were transfected into K562 cells and assayed after 48 hours for luciferase and β-gal expression. We first investigated the activity of each control sequence (ALAS2-1, 2 and 3) containing original or disrupted GATA1 binding site. As predicted and shown previously the sequences with original binding site exhibited more enhancer like activity on luciferase expression than sequence with disrupted (Alt) binding site (Fig 3A) [25].

**Fig 3.**
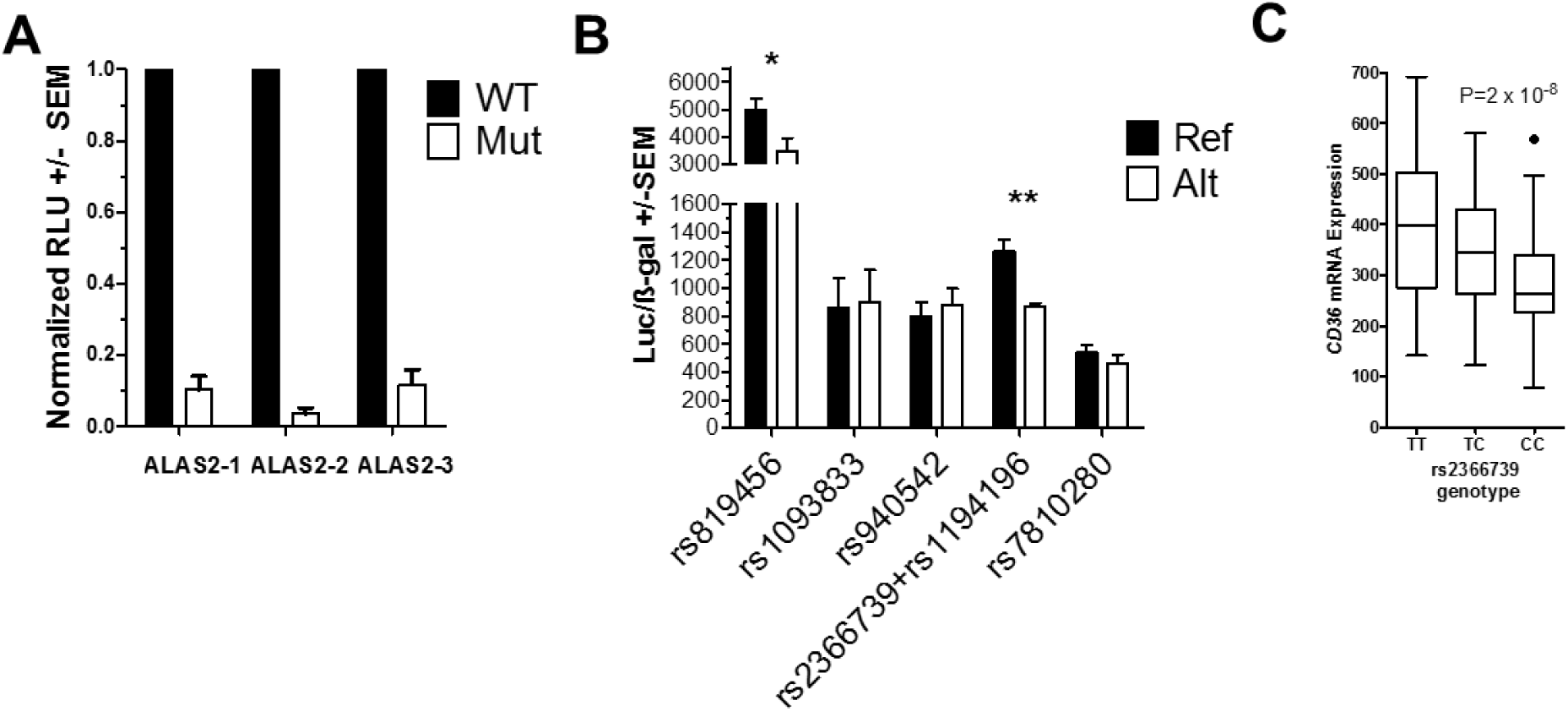
Luciferase Assay of MPRA-identified variants. (A) Relative luciferase activity of reporter vectors containing control (WT) or mutated (Mut) GATA1 sites. (N=3) Significance by one-sample T-test (B) Luciferase activity normalized to β-gal activity. Reference (Ref) and Alternate (Alt) alleles for each variant is indicated. (N=3 to 5) Significance by two-sample T-test. * <0.05, ** < 0.01 (C) Platelet eQTL analysis indicated that platelet *CD36* mRNA levels are associated with rs2366739 genotype (P=2 x 10^−8^). Line = mean expression, Box = 25^th^ to 75 interquartile range (IQR), Whiskers = 1.5 x IQR.

Because eQTLs rs2366739 and rs1194196 are within 21 base pairs of each other and in high linkage disequilibrium we constructed single oligos which contains either reference (T-A for rs2366739-rs1194196) or alternate alleles (C-T) for both eQTLs. These genotypes account for 96% of the observed haplotypes from all populations. Out of the five tested constructs, rs819456 and rs2366739-rs1194196 showed significant transcriptional difference between their reference vs alternate allele as depicted by the luciferase levels (Fig 3B). The rs2366739-rs1194196 results are in agreement with our platelet eQTL data and whole blood eQTL data from Jansen et al. that indicate that the ‘C’ allele of rs2366739 is associated with lower levels of *CD36* mRNA (Fig 3C) [21, 26]. Overall expression from the rs819456 constructs was higher than other constructs but the direction of the difference between alleles (higher in the reference, Fig 3B) was opposite to that of the MPRA results (lower in the reference (Fig 2B)).

rs2366739 has been described as an CD36 eQTL not just in our platelet data but in whole blood data from Võsa et al. (P=1.3 x 10^−267^) [27]. In addition rs2366739 has been associated with DNA methylation levels (P=6.51 x 10^−81^) [28]. These results in addition to the high significance and directional agreement of the rs2366739-rs1194196 luciferase and MPRA results lead us to pursue the rs2366739-rs1194196 locus in further tests.

### Differential protein binding between reference and alternate allele of rs2366739 and rs1194196

One of the mechanisms by which gene expression is regulated is the binding of transcription factors to regulatory elements. To test if the difference in transcription between TA and CT alleles of the rs2366739-rs1194196 constructs is due to alteration in transcription factor binding affinity, we performed an electrophoretic mobility shift assay (EMSA) to compare binding of K562 nuclear extracts to probes derived from the two different haplotypes. The results show formation of a DNA:protein complex with twice as much affinity to the TA genotype probe than to the CT genotype probe (Fig 4). As the luciferase assay suggested that transcription levels are higher for alleles AT, this suggest that the difference in gene transcription could be because be because of higher binding of a transcriptional activator to the AT haplotype.

**Fig 4.**
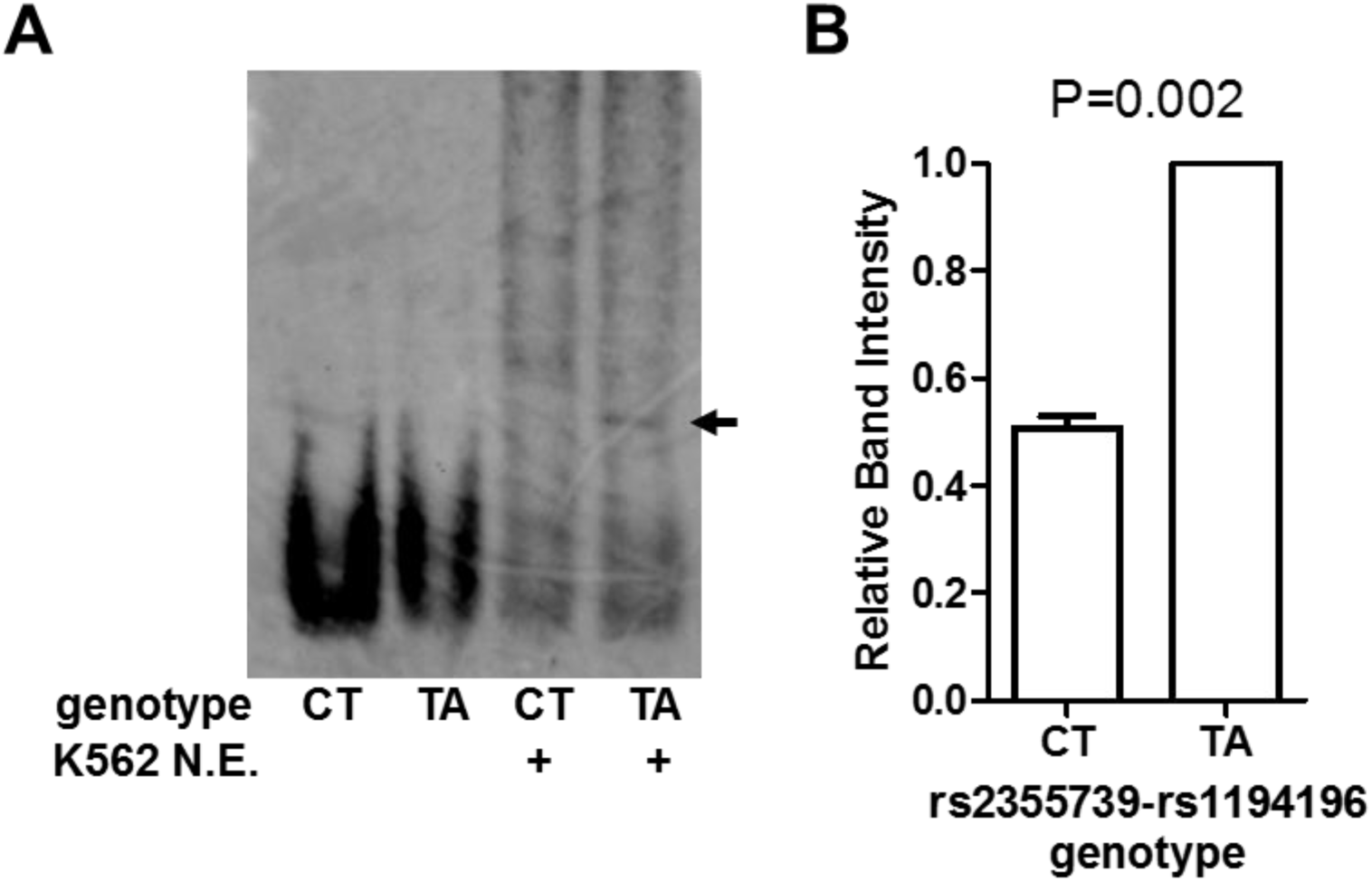
rs2355739-rs1194196 variants alter protein:DNA interactions. (A) Electromobility shift assay using probes containing 70bp of genomic sequence containing SNPs rs2355739 and rs1194196. Digoxin-labeled Probes were incubated with K562 cell nuclear extract where indicated, resolved by gel electrophoresis, and imaged with anti-digoxin antibodies. Arrow indicated specific complex formed by protein:DNA interactions. (B) Quantification of DNA:protein complex by densitometry. (N=3) Significance by one-sample T-test.

### In vivo validation of rs2366739-rs1194196 locus as CD36 regulatory element

To confirm the locus containing rs2366739 and rs1194196 regulates CD36 expression, we generated K562 cell lines with deletion of 573 basepairs containing this region using CRISPR/Cas9. The deletion was confirmed with PCR comparing clones transfected with sgRNAs to those transfected with vectors with no sgRNA (Fig 5, lane C). Clones 4 and 17 did not contain a deletion whereas clones 7 and 20 were successfully altered. To determine the effect of this deletion on *CD36* RNA expression, CD36 transcript levels were measured by qRT-PCR. In the cells with the rs2366739-rs1194196 locus removed (clones 7 and 20), *CD36* mRNA was ∼14 time greater than the clones with the region intact. This supports the evidence that this genomic region identified by MPRA regulated expression of the *CD36* gene.

**Fig 5.**
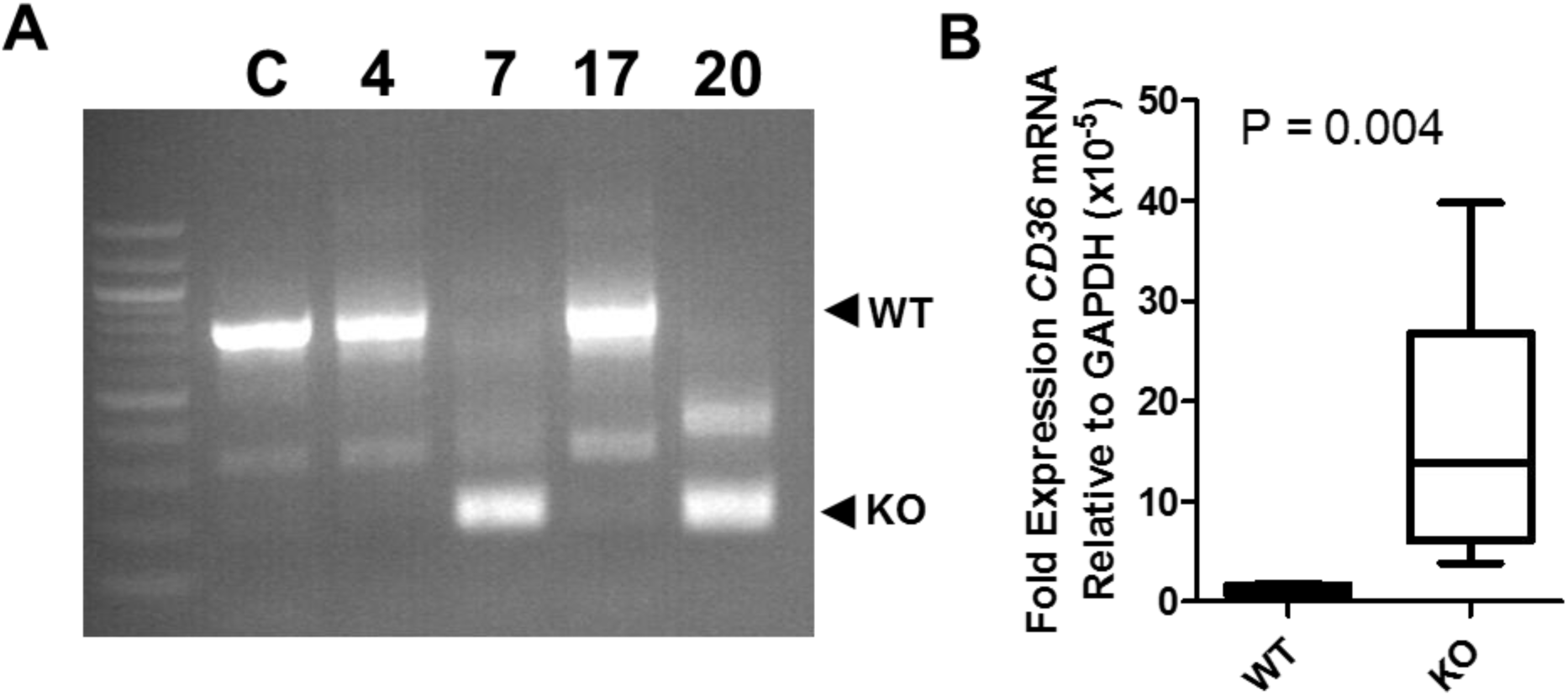
Deletion of the rs2355739-rs1194196 locus increased *CD36* expression. (A) Deletion of the rs2355739-rs1194196 locus by CRISPR/Cas9 was validated by PCR using primers flanking the deleted section. Clones 4 and 17 did not contain a deletion while clones 7 and 20 contained the deletion. (B) qRT-PCR of *CD36* mRNA indicated that the wildtype (WT) clones 4 and 17 contained less *CD36* RNA than clones 7 and 20 lacking the rs2355739-rs1194196 locus (KO).

### Assessment of Clinical Significance

To assess the clinical relevance of our findings we used the PhenoScanner tool, which aggregates large scale GWAS findings. We re-identified the rs2366739 variant in the CD36 associations and we found it to be strongly associated with platelet volume and count but not with phenotypes unrelated to platelet activity. These results reinforce the clinical relevance of our variants and the utility of the MPRA assay.

## Discussion

The expression of CD36 has been actively studied since the identification of a deficiency in healthy Japanese and US donors [29]. CD36 deficiency has been divided into two types: Type I in which neither platelets nor monocytes express surface CD36 and Type II in which only platelets lack expression [30]. Lack of CD36 leads to numerous cellular phenotypes including defective uptake of long chain fatty acids by the myocardium[31, 32] and altered lipid profiles [33, 34]. Lack of CD36 has also been associated with altered foam cell formation in both humans and mice [35, 36]. Even in non-deficient patients a wide range of CD36 expression has been observed and this variation has been associated with the platelet response to oxLDL [19, 37].

Five genetic causes of type I deficiency have been identified, two of which lead to alterations in post-translation modification or surface trafficking, the other three which lead to frame-shift mutations [18, 38-40]. The basis of type II deficiency remains unclear. Mechanisms behind the broad range of non-deficient expression has been explored previously. For example Ghosh et al. previously identified a number of SNPs that are associated with platelet CD36 levels and Masuda explored surface levels in individuals heterozygous for the deficiency mutations mentioned above [19, 37]. However, given the tightly linked nature of the locus (Fig 1), neither of these studies identified functional variants.

Our data presented here represents the first comprehensive analysis and testing of genetic variants associated with *CD36* mRNA expression. We tested 81 variants associated with platelet *CD36* mRNA levels, some of which overlap with variants previously associated with surface levels, by MPRA. Of these variants, rs2366739 and rs1194196, which are within 21 base pairs of each other, were identified by MPRA and validated by (1) altering expression of a linked reporter gene, (2) altering protein binding in a gel-shift assay, and (3) altering endogenous *CD36* expression when the locus containing these variants was deleted by genome editing. Importantly, rs2366739 has previously been associated with both platelet count (P = 9.13 x 10^−10^) and platelet volume (P=5.27 x 10^−11^) [22]. These results narrow down the genomic locus responsible for variable *CD36* expression from ∼125kb to 21bp, making identification of the mechanism feasible. This would not be possible with GWAS or eQTL analysis alone.

The analysis methods employed for our CD36 MPRA are another important contribution of this work. We applied t- and U-tests that have been employed in prior MPRA studies, but we also introduced a new and more comprehensive Bayesian approach. Our new method recapitulates the findings of the prior methods, but also identifies a variant that was missed by the prior approaches. A key advantage of our new statistical method is a generative stochastic model that probabilistically accounts for the discrete nature of NGS count data as well as sources of variation in the MPRA experiment without having to discard zero counts. Fig S6 shows a Kruschke diagram of the generative process considered by the model. The sources of variation addressed include variation in the barcode abundances in the cDNA library as well as other factors. Furthermore, the use of empirical priors provides estimate shrinkage; noisy parameter estimates are shrunken towards more moderate levels observed throughout the rest of the assay. This process helps eliminate false positives without the heavy statistical burden of multiple testing correction procedures like false discovery rates. We have previously studied the statistical power of MPRA experiments using the standard approaches [24]. Although the prior methods were at least partially successful to analyze MPRA data, our Bayesian model appears both practical and more powerful. Therefore, our paper demonstrates effective statistical improvements to analysis of MPRA studies. This approach may be particularly important in larger scale genome wide MPRA studies, and more work is warranted to improve the cost-effectiveness as well as the discovery potential of MPRA assays.

One of the weaknesses of this study is that we have been unable to identify the DNA-binding protein factor whose binding is altered by the variants. The rs2366739-rs1194196 is located in a genomic locus that contains no strong annotation signals in the ENCODE, BloodChIP, or BLUEPRINT databases [41-43]. However, this locus is contained within a THE1B repeated element. THE1B is a mammalian apparent LTR retrotransposon. A reactivated THE1B element has previously been reported to lead to activation of the *CSF1R* proto-oncogene in human lymphoma [44]. It is possible that the variants cause alteration of the transcription factor binding sites contained within this element, leading to altered *CD36* expression. More work is needed to determine if the rs2366739-rs1194196 locus is also responsible for regulating expression in other cell types, such as monocytes.

Currently, more human genetic studies are moving beyond associations of genotypes with phenotypes to seeking the molecular mechanisms behind the variants responsible for the observed traits. Mechanistic understanding of genetic variants will provide better understanding of the observed physiology, allow for more precise biomarkers, and identify potential new therapeutic targets. Given the large number of variants potentially associated with a trait in highly linked genomic regions such as *CD36*, high-throughput methods are necessary to efficiently test and identify functional polymorphisms in an unbiased manner. Our identification of the genetic locus responsible for inter-individual variation in non-deficient *CD36* expression opens new areas of investigation into the link between this locus and platelet function, serum lipid levels, and atherosclerosis.

## Materials and Methods

### Cell culture

K562 Cells from ATCC (CCL-243) were maintained in RPMI 1640 media (Invitrogen, CA 10-040-cv) containing penicillin-streptomycin and 10% FBS.

### MPRA

The MPRA design was based on the method previously published [45]. To design the *CD36* MPRA library, we used a p-value threshold of p < 1×10^−6^ to select the expression quantitative trait loci (eQTLs) most highly associated with *CD36* expression in the PRAX study surrounded by 150bp of hg38 genomic context [21]. After discarding 5 eQTLs that contained digestion sites for restriction enzymes used in the library preparation in their genomic context, this yielded a set of 81 CD36 SNPs to assay. We also included 10 SNPs previously identified in the literature as directly affecting *ALAS2, HBG2, PKRRE*, and *URUOS* expression as a set of positive controls (Table S1) [46, 47]. We synthesized 40 oligonucleotide replicates per allele(Agilent, Santa Clara, CA), each uniquely tagged with inert 10bp barcodes which followed the design criteria stipulated in Melnikov et al. [48]. The library of oligonucleotides was amplified using emulsion PCR and the primers 5’-TGCTAAGGCCTAACTGGCCAG-3’ and 5’-CTCGGCGGCCAAGTATTCAT-3’ which also added additional sequence containing SfiI restriction enzyme sites to each end. The design of the oligonucleotide library is shown in Fig S1. After each oligo was directionally cloned into pMPRA1 (Addgene, Cambridge, MA) using the SfiI sites, a minimal promoter and luciferase cassette derived from pNL3.2 (Promega, Madison, WI) was inserted in between the genomic sequences and the barcode using KpnI and XbaI sites. This library of plasmids was transfected into K562 cells and then RNA was harvested 48 hours later. After reverse transcription with a polyT primer, three separate amplifications of the cDNA were performed to generate RNA sequencing libraries. The barcodes contained in the MPRA plasmid library were subjected to two separate amplifications to generate DNA sequencing libraries. RNA and plasmid barcode expression was quantified by next generation sequencing on an Illumina MiSeq in the Children’s Hospital of Philadelphia sequencing core. After extraction from the fastq files, barcodes with a quality of Q>30 at every base was counted. RNA barcode counts were analyzed in conjunction with the DNA barcode counts to control for variances in barcode abundances introduced by library generation. Quality control data concerning the sequencing libraries are presented in Table S2 and Figs S2-S3.

Both frequentist (t-tests and U-tests) and Bayesian analyses were applied to identify variants that caused alterations to transcription activity by use of RNA-to-DNA ratios to examine allelic differences. Analysis of the barcode counts proceeds by computing the MPRA activity of each barcode as the log-ratio of the depth-normalized RNA counts to depth-normalized DNA counts and each variant’s corresponding “transcription shift”. The transcription shift of a variant is defined as the difference in activity between the alternate and reference alleles.

The application of traditional frequentist tests to MPRA activities has been previously described [24, 25]. The novel Bayesian analysis models the barcode count data using negative binomial distributions with empirical gamma priors (Figs S4), this analysis method provides greater sensitivity than traditional methods while retaining specificity.

#### Luciferase assay

We designed and ordered oligonucleotides (IDT-DNA, Coralville, IA) containing the candidate MPRA-functional SNPs and 40bp of flanking genomic sequence (Table S3). The amplified sequence was inserted into nano-luciferase containing plasmid pNL3.2 (Promega, Madison, WI) using HindIII and XhoI (ThermoFisher, Waltham, MA) restriction sites. The sequence was confirmed by sequencing. The reporter plasmids and β-gal expression plasmids were cotransfected in K562 cells and luciferase assay was carried out after 48 hours using Nanoglo luciferase kit (Promega) and normalized to ß-gal expression measured using assay reagent (ThermoFisher).

#### Electromobility assay (EMSA)

Nuclear extracts from K562 cells were isolated using the NE-PER Nuclear and Cytoplasmic Extraction Kit (ThermoFisher). 70 base pair sequences containing either the reference alleles for rs2366739 and rs1194196 (referred to as TA) or alternate alleles (referred to as CT) were amplified by PCR using MPRA plasmid library as template. Digoxigenin (DIG) labeled nucleotides (Roche, Basel, Switzerland) were used to create amplified sequences with DIG labeled base pairs. The sequences were purified by agarose gel electrophoresis and the QIAEX II Gel Extraction Kit (Qiagen, Hilden, Germany).

10μg of nuclear extract was incubated with DIG labeled probes in buffer containing 10% glycerol, 20 mM HEPES, 30 mM KCl, 30 mM NaCl, 3 mM MgCl_2_, 1 mM DTT. 1 μg of Poly dI-dC was added to reduce nonspecific binding. The reaction was carried out for 30 mins on ice. The sample was mixed with 10X orange loading dye (Licor, Lincoln, NE) and loaded on 6% acrylamide gel and ran for 4-5 hours with 0.5% TBE buffer. The DNA-protein complex was transferred on positively charged Biodyne nylon membrane (Pall Industries, Fort Washington, NY) using 0.5% TBE for 45 minutes. The membrane was incubated in blocking solution for 30 mins at RT, followed by Anti-Digoxigenin-AP antibody containing blocking solution for 30 mins at RT. The membrane was then washed twice for 15 minutes each using washing solution, visualized using chemiluminescence, and quantified.

#### CRISPR-modified cell line generation

To generate genomic deletion mutants, the CRISPR/Cas9 system was used as previously reported [49]. Two guide RNAs (sgRNA) flanking 573 base pairs containing rs2366739 and rs1194196 were designed (IDT DNA). The design of sgRNA pairs for targeting and prediction of off-target sites were based on online tools: CRISPR Design (http://crispr.mit.edu/) and CRISPOR (http://www.crispor.org)[50]. Two guide RNAs, 5’-TACCCCCATTGTATCTATCTAGG-3’ and 5’-CTACAGTAAATACACTTGTCAGG -3’ were used to delete the 573 basepair region.

Pairs of complementary DNA oligos (IDTDNA, Coralville, IA) were individually phosphorylated with T4 polynucleotide kinase (NEB, Ipswitch, MA) and then annealed. Each DNA oligo duplex had 5’ overhangs (forward: ACCG, reverse: AAAC) designed to be directly cloned into the BbsI or BsaI-digested and dephosphorylated AIO-GFP(Cas9) vector using the Quick Ligation Kit (NEB). The first and second sgRNA was cloned into the BbsI and BsaI sites, respectively, and confirmed by colony PCR and sequencing. The plasmid was transfected in K562 using Lipofectamine2000 (ThermoFisher). After 24 hours, the GFP positive cells were sorted by flow cytometry and individually seeded in a 96 well plate. Single colonies were expanded further and a cell line was established from a single clone. Deletion was confirmed by PCR and sequencing.

#### Reverse transcription and quantitative real time PCR

Total RNA was isolated using Trizol Reagent (ThermoFisher). 3μg total RNA was used for first strand cDNA synthesis with the SuperScript™ III First-Strand Synthesis System (ThermoFisher). To evaluate relative expression levels of mRNAs, we performed qRT-PCR with the Power SYBR Green PCR master mix (Life Technologies, Carlsbad, CA) normalized to Actin. We carried out real time PCR reaction and analyses in 384-well optical reaction plates using the CFX384 instrument (Bio-Rad, Hercules, CA).

## Acknowledgements

This work was supported by National Institutes of Health – National Heart, Lung and Blood Institute grant R01HL128234 and the Cardeza Foundation for Hematologic Research. We also thank the members of the Children’s Hospital of Philadelphia Nucleic Acid PCR Research Core Facility for help in preparation of sequencing libraries.

