## Supplemental File for "Identification of functional variants for platelet *CD36* expression by Massively Parallel Reporter Assay"

**Table S1: Sequences of MPRA Controls**

| Gene | Type | Sequence |
|---|---|---|
| PKRRE | Ref | AGGAAATGCCAGGAGATGAGGGCAGAGAGCAGGCCGTTCTGGGGGAGGGATTCTGTGGGGACAGGGTGGCCTACTGGGTGTGCCCCTTTTCTCTTCTCT GTCTCCCTTAGATAAGACCAGCAGTTTTGTCATCCTCTCCCTCTCATTCCA |
| PKRRE | Mut | AGGAAATGCCAGGAGATGAGGGCAGAGAGCAGGCCGTTCTGGGGGAGGGATTCTGTGGGGACAGGGTGGCCTACTGGGTGTGCCCCTTTTCTCTTATCT GTCTCCCTTAGATAAGACCAGCAGTTTTGTCATCCTCTCCCTCTCATTCCA |
| PKRRE | Ref | AGGAAATGCCAGGAGATGAGGGCAGAGAGCAGGCCGTTCTGGGGGAGGGATTCTGTGGGGACAGGGTGGCCTACTGGGTGTGCCCCTTTTCTCTTCTCT GTCTCCCTTAGATAAGACCAGCAGTTTTGTCATCCTCTCCCTCTCATTCCA |
| PKRRE | Mut | AGGAAATGCCAGGAGATGAGGGCAGAGAGCAGGCCGTTCTGGGGGAGGGATTCTGTGGGGACAGGGTGGCCTACTGGGTGTGCCCCTTTTCTCTTCGCT GTCTCCCTTAGATAAGACCAGCAGTTTTGTCATCCTCTCCCTCTCATTCCA |
| PKRRE | Ref | AGGAAATGCCAGGAGATGAGGGCAGAGAGCAGGCCGTTCTGGGGGAGGGATTCTGTGGGGACAGGGTGGCCTACTGGGTGTGCCCCTTTTCTCTTCTCT GTCTCCCTTAGATAAGACCAGCAGTTTTGTCATCCTCTCCCTCTCATTCCA |
| PKRRE | Mut | AGGAAATGCCAGGAGATGAGGGCAGAGAGCAGGCCGTTCTGGGGGAGGGATTCTGTGGGGACAGGGTGGCCTACTGGGTGTGCCCCTTTTCTCTTCTAT GTCTCCCTTAGATAAGACCAGCAGTTTTGTCATCCTCTCCCTCTCATTCCA |
| PKRRE | Ref | AGGAAATGCCAGGAGATGAGGGCAGAGAGCAGGCCGTTCTGGGGGAGGGATTCTGTGGGGACAGGGTGGCCTACTGGGTGTGCCCCTTTTCTCTTCTCT GTCTCCCTTAGATAAGACCAGCAGTTTTGTCATCCTCTCCCTCTCATTCCA |
| PKRRE | Mut | AGGAAATGCCAGGAGATGAGGGCAGAGAGCAGGCCGTTCTGGGGGAGGGATTCTGTGGGGACAGGGTGGCCTACTGGGTGTGCCCCTTTTCTCTTCTCG GTCTCCCTTAGATAAGACCAGCAGTTTTGTCATCCTCTCCCTCTCATTCCA |
| PKRRE | Ref | AGGAAATGCCAGGAGATGAGGGCAGAGAGCAGGCCGTTCTGGGGGAGGGATTCTGTGGGGACAGGGTGGCCTACTGGGTGTGCCCCTTTTCTCTTCTCT GTCTCCCTTAGATAAGACCAGCAGTTTTGTCATCCTCTCCCTCTCATTCCA |
| PKRRE | Mut | AGGAAATGCCAGGAGATGAGGGCAGAGAGCAGGCCGTTCTGGGGGAGGGATTCTGTGGGGACAGGGTGGCCTACTGGGTGTGCCCCTTTTCTCTTCTCT CTCTCCCTTAGATAAGACCAGCAGTTTTGTCATCCTCTCCCTCTCATTCCA |
| ALAS2 | Ref | CTGGAGATATGTGTTTCCCTTCCCCTGCCTGCTTGTGAAAGCTAAAGCACTTGGGGCTGAGCCTGCAGACCACAGATAAAGTTGCCAGAGTTTATCGCCATT GGGGTCTGACCACTCCCCAAGCTGAGACCAGGGCGTGGGAGAGAAGAG |
| ALAS2 | Mut | CTGGAGATATGTGTTTCCCTTCCCCTGCCTGCTTGTGAAAGCTAAAGCACTTGGGGCTGAGCCTGCAGACCACACATAAAGTTGCCAGAGTTTATCGCCATT GGGGTCTGACCACTCCCCAAGCTGAGACCAGGGCGTGGGAGAGAAGAG |
| ALAS2 | Ref | TGGAGATATGTGTTTCCCTTCCCCTGCCTGCTTGTGAAAGCTAAAGCACTTGGGGCTGAGCCTGCAGACCACAGATAAAGTTGCCAGAGTTTATCGCCATT GGGGTCTGACCACTCCCCAAGCTGAGACCAGGGCGTGGGAGAGAAGAGG |
| ALAS2 | Mut | TGGAGATATGTGTTTCCCTTCCCCTGCCTGCTTGTGAAAGCTAAAGCACTTGGGGCTGAGCCTGCAGACCACAGGTAAAGTTGCCAGAGTTTATCGCCATT GGGGTCTGACCACTCCCCAAGCTGAGACCAGGGCGTGGGAGAGAAGAGG |
| ALAS2 | Ref | GGAGATATGTGTTTCCCTTCCCCTGCCTGCTTGTGAAAGCTAAAGCACTTGGGGCTGAGCCTGCAGACCACAGATAAAGTTGCCAGAGTTTATCGCCATTG GGGTCTGACCACTCCCCAAGCTGAGACCAGGGCGTGGGAGAGAAGAGGA |
| ALAS2 | Mut | GGAGATATGTGTTTCCCTTCCCCTGCCTGCTTGTGAAAGCTAAAGCACTTGGGGCTGAGCCTGCAGACCACAGACAAAGTTGCCAGAGTTTATCGCCATTG GGGTCTGACCACTCCCCAAGCTGAGACCAGGGCGTGGGAGAGAAGAGGA |
| UROS | Ref | TGGAAGAGACCTATCCCTCCTACACCATGACTTAGCACTAATGGGCTTGTTCTTTCTGAAGACCCCTGTCACTGATAAGGCCAAGAAAGAGCATGTTAGCA GTTGATATCACTTGGAAAGACAGATGAAACCATTAGTGAAAGCCAAGTG |

| | | |
|---|---|---|
| UROS | Mut | TGGAAGAGACCTATCCCTCCTACACCATGACTTAGCACTAATGGGCTTGTTCTTTCTGAAGACCCCTGTCACTGGTAAGGCCAAGAAAGAGCATGTTAGCAGTTGATATCACTTGGAAAGACAGATGAAACCATTAGTGAAAGCCAAGTG |
| SPI1 | Ref | GTTGTTTAAAATTTTCTCTGAGACACGACGTTCTCCTACCTAACTCTTATTGAAATGAACAAAAAACCCTCCTCCAATTACCCACCCTCCCCACTCTCCCCTCTAGACTAAATGTATGCCTGAACTCTGAGGGTTCCGTCATGGGGCCCA |
| SPI1 | Mut | GTTGTTTAAAATTTTCTCTGAGACACGACGTTCTCCTACCTAACTCTTATTGAAATGAACAAAAAACCCTCCTCTAATTACCCACCCTCCCCACTCTCCCCTCTAGACTAAATGTATGCCTGAACTCTGAGGGTTCCGTCATGGGGCCCA |
| HBG2 | Ref | TAACATTAATCTATTCCTGCACTGAAACTGTTGCTTTATAGGATTTTTCACTACACTAATGAGAACTTAAGAGATAATGGCCTAAAACCACAGAGAGTATATTCAAAGATAAGTATAGCACTTCTTATTTGGAAACCAATGCTTACTAAA |
| HBG2 | Mut | TAACATTAATCTATTCCTGCACTGAAACTGTTGCTTTATAGGATTTTTCACTACACTAATGAGAACTTAAGAGAGAATGGCCTAAAACCACAGAGAGTATATTCAAAGATAAGTATAGCACTTCTTATTTGGAAACCAATGCTTACTAAA |

**Table S2: RNA-Seq Quality**

| Sample | Sample Depth | Number of Reads | Fraction Usable Reads | Fraction 0-count barcodes |
|---|---|---|---|---|
| cDNA1 | 1891321 | 2739571 | 0.69 | 0.09 |
| cDNA2 | 2967012 | 4109045 | 0.72 | 0.09 |
| cDNA3 | 1577854 | 2242279 | 0.70 | 0.12 |
| Plasmid1 | 1968644 | 2757765 | 0.71 | 0.07 |
| Plasmid2 | 3023219 | 4342543 | 0.70 | 0.06 |

**Table S3.** Sequences generated for each variant and for luciferase assays.

| SNP-genotype | Sequence |
|---|---|
| rs1093833 T (Ref) | GCAGCAGCCACAACCAAGTGTTAGTGGGTATAGGGGTGCC |
| rs1093833 C (Alt) | GCAGCAGCCACAACCAAGTGCTAGTGGGTATAGGGGTGCC |
| rs7810280 G (Ref) | AATCAACCAAGATGATTTACGGGTCAAAGTATTATAGTGT |
| rs7810280 A (Alt) | AATCAACCAAGATGATTTACAGGTCAAAGTATTATAGTGT |
| rs819456 T (Ref) | TGCTGTTCTTGTAATAGTGATTGGGTCTCATGAAATCTGA |
| rs819456 A (Alt) | TGCTGTTCTTGTAATAGTGAATGGGTCTCATGAAATCTGA |
| rs940542T (Ref) | TTTAACTGAATTTTTAATGTTGTTAACTGAGATAAGTGAA |
| Rs940542 C (Alt) | TTTAACTGAATTTTTAATGTCGTTAACTGAGATAAGTGAA |
| rs2366739+rs1194196 TA (Ref) | TCCCATGCTGTTCTTGTAATAGTGATTGGGTCTCATGAAATCTGATGTTTTTAAAAACGG GAGTTTCTCTGAACAGTCTC |
| rs2366739+rs1194196 CT (Alt) | TCCCATGCTGTTCTTGTAATAGTGACTGGGTCTCATGAAATCTGATGTTTTTATAAACGG GAGTTTCTCTGAACAGTCTC |

**Table S4. Complete MPRA results of positive Controls**

| Control | Transcription Shift | t-test | | u-test | | Bayesian Model | |
|---|---|---|---|---|---|---|---|
| | | P-value | Q-value | P-value | Q-value | Posterior Mean | 95% Credible Interval |
| URUOS | -2.55525 | 1.94E-131 | 1.76E-129 | 2.08E-38 | 1.89E-36 | -2.44298 | -2.616 : -2.267 |
| PKRRE5 | -1.00354 | 9.86E-73 | 4.49E-71 | 1.41E-37 | 6.40E-36 | -1.11463 | -1.282 : -0.944 |
| PKRRE3 | -0.87213 | 1.24E-71 | 3.77E-70 | 4.44E-37 | 1.35E-35 | -1.01771 | -1.183 : -0.848 |
| ALAS23 | -2.08592 | 8.95E-62 | 2.04E-60 | 5.82E-33 | 8.82E-32 | -2.00429 | -2.276 : -1.703 |
| ALAS22 | -2.413 | 2.53E-57 | 3.83E-56 | 1.11E-28 | 1.12E-27 | -2.48037 | -2.838 : -2.114 |
| ALAS21 | -2.10623 | 6.23E-57 | 8.10E-56 | 1.52E-32 | 1.97E-31 | -2.00922 | -2.33 : -1.699 |
| PKRRE4 | -0.89499 | 2.37E-56 | 2.70E-55 | 7.73E-36 | 1.76E-34 | -0.93448 | -1.097 : -0.769 |
| PKRRE1 | -0.69606 | 7.38E-39 | 7.46E-38 | 2.57E-30 | 2.92E-29 | -0.75478 | -0.924 : -0.59 |
| PKRRE2 | -0.26871 | 1.76E-06 | 1.45E-05 | 1.47E-08 | 1.34E-07 | -0.37922 | -0.698 : -0.055 |
| HBG2 | 0.06142 | 4.96E-01 | 7.28E-01 | 7.87E-01 | 8.97E-01 | -0.01277 | -0.231 : 0.214 |

**Table S5.** Complete Results of MPRA of CD36 variants.

| Control | Transcription Shift | t-test | | u-test | | Bayesian Model | |
|---|---|---|---|---|---|---|---|
| | | P-value | Q-value | P-value | Q-value | Posterior Mean | 95% Credible Interval |
| rs2366739 | -1.04532 | 6.87E-59 | 1.25E-57 | 1.41E-35 | 2.57E-34 | -1.13049 | -1.295 : -0.955 |
| rs940542 | -0.2785 | 2.56E-07 | 2.33E-06 | 6.51E-07 | 5.39E-06 | -0.29285 | -0.481 : -0.12 |
| rs1093831 | -0.273 | 1.07E-04 | 8.09E-04 | 7.70E-05 | 5.84E-04 | 0.10111 | -0.113 : 0.315 |
| rs6467258 | 0.22024 | 6.72E-03 | 4.37E-02 | 7.47E-04 | 5.23E-03 | -0.05448 | -0.259 : 0.16 |
| rs1194196 | -0.23346 | 1.21E-02 | 6.45E-02 | 1.07E-03 | 6.93E-03 | -0.08582 | -0.271 : 0.111 |
| rs11464747 | 0.36056 | 1.02E-03 | 7.16E-03 | 1.70E-03 | 1.03E-02 | 1.23996 | 0.962 : 1.518 |
| rs819456 | 0.31541 | 3.05E-02 | 1.32E-01 | 2.68E-03 | 1.52E-02 | 0.46373 | 0.079 : 0.861 |
| rs6961069 | 0.21993 | 1.14E-02 | 6.45E-02 | 2.90E-03 | 1.55E-02 | 0.01234 | -0.211 : 0.233 |
| rs819457 | 0.18773 | 1.58E-02 | 7.58E-02 | 3.07E-03 | 1.55E-02 | 0.14833 | -0.072 : 0.372 |
| rs28851188 | 0.20479 | 9.63E-03 | 5.84E-02 | 1.53E-02 | 6.94E-02 | 0.07836 | -0.132 : 0.282 |
| rs1761661 | -0.48586 | 1.51E-01 | 3.70E-01 | 1.47E-02 | 6.94E-02 | -0.44408 | -1.109 : 0.227 |
| rs1608671 | 0.2257 | 1.32E-02 | 6.69E-02 | 1.83E-02 | 7.91E-02 | 0.24244 | -0.001 : 0.463 |
| rs13233631 | 0.23128 | 6.44E-02 | 2.17E-01 | 2.29E-02 | 9.47E-02 | -0.03622 | -0.305 : 0.253 |
| rs819442 | -0.14388 | 1.59E-01 | 3.77E-01 | 2.45E-02 | 9.67E-02 | 0.05905 | -0.24 : 0.361 |
| rs819445 | -0.01314 | 9.05E-01 | 9.89E-01 | 3.11E-02 | 1.18E-01 | -0.02339 | -0.289 : 0.237 |
| rs1761667 | -0.18177 | 4.86E-02 | 1.84E-01 | 3.85E-02 | 1.40E-01 | -0.08644 | -0.315 : 0.148 |

| | | | | | | | |
|---|---|---|---|---|---|---|---|
| rs4626520 | 0.13624 | 6.17E-02 | 2.17E-01 | 4.96E-02 | 1.67E-01 | 0.02326 | -0.188 : 0.223 |
| rs2781841 | -0.1183 | 1.50E-01 | 3.70E-01 | 4.85E-02 | 1.67E-01 | -0.10839 | -0.332 : 0.128 |
| rs1851935 | 0.46972 | 2.08E-02 | 9.44E-02 | 5.60E-02 | 1.82E-01 | 0.58079 | 0.15 : 0.982 |
| rs2177616 | -0.09018 | 1.68E-01 | 3.82E-01 | 6.19E-02 | 1.94E-01 | -0.00484 | -0.188 : 0.191 |
| rs1194197 | 0.11786 | 1.07E-01 | 3.03E-01 | 7.32E-02 | 2.08E-01 | 0.12638 | -0.085 : 0.338 |
| rs4731643 | -0.15098 | 1.11E-01 | 3.05E-01 | 7.11E-02 | 2.08E-01 | -0.11007 | -0.36 : 0.128 |
| rs7810280 | -0.11062 | 3.32E-01 | 6.05E-01 | 7.19E-02 | 2.08E-01 | -0.22907 | -0.463 : 0.009 |
| rs10233710 | 0.16293 | 3.71E-02 | 1.53E-01 | 8.50E-02 | 2.34E-01 | 0.26916 | 0.051 : 0.485 |
| rs1194179 | 0.05632 | 4.24E-01 | 6.88E-01 | 9.44E-02 | 2.53E-01 | -0.14227 | -0.349 : 0.06 |
| rs1953298 | -0.07658 | 5.12E-01 | 7.39E-01 | 1.13E-01 | 2.95E-01 | -0.0912 | -0.372 : 0.176 |
| rs1093833 | 0.54135 | 1.04E-01 | 3.03E-01 | 1.23E-01 | 3.12E-01 | 1.00913 | 0.271 : 1.764 |
| rs2366855 | -0.04748 | 6.02E-01 | 8.05E-01 | 1.28E-01 | 3.15E-01 | -0.26947 | -0.505 : -0.034 |
| rs12155030 | 0.49526 | 3.87E-02 | 1.53E-01 | 1.43E-01 | 3.24E-01 | 0.21049 | -0.404 : 0.784 |
| rs3211821 | 0.15352 | 6.21E-02 | 2.17E-01 | 1.40E-01 | 3.24E-01 | 0.17309 | -0.052 : 0.38 |
| rs138369160 | 0.1948 | 8.07E-02 | 2.62E-01 | 1.46E-01 | 3.24E-01 | 0.31868 | 0.018 : 0.617 |
| rs17154155 | -0.13435 | 8.45E-02 | 2.65E-01 | 1.37E-01 | 3.24E-01 | -0.23316 | -0.449 : -0.009 |
| rs4728183 | 0.00832 | 9.30E-01 | 9.89E-01 | 1.50E-01 | 3.24E-01 | 0.01875 | -0.223 : 0.266 |
| rs34592988 | 0.07016 | 4.06E-01 | 6.73E-01 | 1.55E-01 | 3.29E-01 | 0.0955 | -0.119 : 0.308 |
| rs819436 | 0.11231 | 3.14E-01 | 5.95E-01 | 1.75E-01 | 3.61E-01 | 0.16837 | -0.111 : 0.458 |
| rs9649532 | 0.12441 | 3.66E-01 | 6.28E-01 | 1.94E-01 | 3.92E-01 | 0.0279 | -0.34 : 0.4 |

| | | | | | | | |
|---|---|---|---|---|---|---|---|
| rs3211842 | 0.09516 | 1.32E-01 | 3.55E-01 | 2.17E-01 | 4.21E-01 | 0.24792 | 0.042 : 0.445 |
| rs1194177 | -0.01999 | 8.35E-01 | 9.75E-01 | 2.16E-01 | 4.21E-01 | -0.33384 | -0.573 : -0.086 |
| rs28854232 | -0.01207 | 8.93E-01 | 9.89E-01 | 2.24E-01 | 4.25E-01 | -0.06846 | -0.295 : 0.152 |
| rs1761662 | -0.10311 | 2.55E-01 | 5.51E-01 | 2.31E-01 | 4.28E-01 | -0.0865 | -0.33 : 0.161 |
| rs1049654 | -0.12294 | 1.62E-01 | 3.77E-01 | 2.72E-01 | 4.88E-01 | -0.07185 | -0.299 : 0.16 |
| rs1851934 | 0.09629 | 4.03E-01 | 6.73E-01 | 2.73E-01 | 4.88E-01 | 0.40852 | 0.138 : 0.684 |
| rs1320408 | 0.06631 | 2.66E-01 | 5.62E-01 | 2.85E-01 | 4.90E-01 | -0.0141 | -0.207 : 0.173 |
| rs1761645 | -0.02737 | 7.51E-01 | 9.14E-01 | 2.85E-01 | 4.90E-01 | -0.23063 | -0.459 : -0.012 |
| rs1093829 | 0.12957 | 1.48E-01 | 3.70E-01 | 3.23E-01 | 5.16E-01 | 0.11022 | -0.115 : 0.35 |
| rs1761646 | 0.08586 | 2.86E-01 | 5.75E-01 | 3.10E-01 | 5.16E-01 | 0.14086 | -0.064 : 0.371 |
| rs4316098 | 0.07598 | 4.62E-01 | 7.08E-01 | 3.19E-01 | 5.16E-01 | 0.10976 | -0.145 : 0.356 |
| rs1953299 | 0.0382 | 6.67E-01 | 8.74E-01 | 3.21E-01 | 5.16E-01 | -0.5241 | -0.742 : -0.308 |
| rs9649527 | 0.06096 | 4.33E-01 | 6.92E-01 | 3.34E-01 | 5.24E-01 | -0.23275 | -0.485 : 0.014 |
| rs11770358 | 0.15989 | 2.91E-01 | 5.75E-01 | 3.44E-01 | 5.31E-01 | 0.13013 | -0.193 : 0.463 |
| rs1093830 | -0.00965 | 8.35E-01 | 9.75E-01 | 3.65E-01 | 5.53E-01 | -0.0652 | -0.239 : 0.118 |
| rs819460 | 0.14181 | 2.72E-01 | 5.63E-01 | 3.72E-01 | 5.55E-01 | 0.15343 | -0.127 : 0.427 |
| rs1194182 | -0.2146 | 9.11E-02 | 2.76E-01 | 4.08E-01 | 5.89E-01 | -0.33051 | -0.6 : -0.05 |
| rs1761673 | 0.03448 | 7.63E-01 | 9.14E-01 | 4.08E-01 | 5.89E-01 | 0.11188 | -0.133 : 0.356 |
| rs7793698 | 0.04443 | 7.36E-01 | 9.14E-01 | 4.29E-01 | 6.09E-01 | -0.02528 | -0.334 : 0.286 |
| rs13236689 | 0.09468 | 3.04E-01 | 5.89E-01 | 4.91E-01 | 6.87E-01 | 0.03031 | -0.207 : 0.262 |

| | | | | | | | |
|---|---|---|---|---|---|---|---|
| rs1093834 | -0.00567 | 9.20E-01 | 9.89E-01 | 5.05E-01 | 6.96E-01 | 0.15143 | -0.051 : 0.35 |
| rs6467251 | 0.17471 | 4.67E-01 | 7.08E-01 | 6.15E-01 | 8.36E-01 | -0.03797 | -0.526 : 0.432 |
| rs701269 | 0.08099 | 3.24E-01 | 6.01E-01 | 6.55E-01 | 8.46E-01 | -0.28239 | -0.522 : -0.045 |
| rs34736275 | -0.02584 | 6.72E-01 | 8.74E-01 | 6.53E-01 | 8.46E-01 | 0.02446 | -0.157 : 0.212 |
| rs819455 | 0.00715 | 9.37E-01 | 9.89E-01 | 6.60E-01 | 8.46E-01 | 0.00605 | -0.23 : 0.231 |
| rs7794010 | -0.00445 | 9.53E-01 | 9.89E-01 | 6.53E-01 | 8.46E-01 | 0.24392 | 0.018 : 0.468 |
| rs1527479 | 0.15073 | 2.47E-01 | 5.49E-01 | 7.24E-01 | 8.71E-01 | 0.13982 | -0.109 : 0.389 |
| rs12706912 | 0.10766 | 3.43E-01 | 6.12E-01 | 6.97E-01 | 8.71E-01 | -0.08802 | -0.336 : 0.168 |
| rs1537477 | 0.02838 | 5.91E-01 | 8.02E-01 | 7.37E-01 | 8.71E-01 | 0.05378 | -0.139 : 0.237 |
| rs1194178 | 0.02119 | 8.72E-01 | 9.89E-01 | 7.25E-01 | 8.71E-01 | 0.15633 | -0.149 : 0.443 |
| rs1054516 | -0.0088 | 8.97E-01 | 9.89E-01 | 7.12E-01 | 8.71E-01 | 0.01406 | -0.188 : 0.212 |
| rs2366744 | -0.00327 | 9.57E-01 | 9.89E-01 | 7.36E-01 | 8.71E-01 | -0.09881 | -0.285 : 0.093 |
| rs819444 | 0.03194 | 6.87E-01 | 8.80E-01 | 7.55E-01 | 8.81E-01 | 0.02406 | -0.266 : 0.319 |
| rs1194195 | -0.082 | 4.59E-01 | 7.08E-01 | 8.18E-01 | 8.97E-01 | -0.11836 | -0.359 : 0.126 |
| rs1093835 | -0.03549 | 5.52E-01 | 7.64E-01 | 8.02E-01 | 8.97E-01 | -0.0405 | -0.22 : 0.134 |
| rs3212160 | 0.04682 | 5.54E-01 | 7.64E-01 | 8.10E-01 | 8.97E-01 | 0.01562 | -0.196 : 0.232 |
| rs819443 | -0.00145 | 9.89E-01 | 9.89E-01 | 8.14E-01 | 8.97E-01 | -0.09974 | -0.433 : 0.226 |
| rs6961024 | 0.0029 | 9.72E-01 | 9.89E-01 | 8.49E-01 | 9.19E-01 | 0.26364 | 0.068 : 0.472 |
| rs10499859 | 0.09846 | 3.50E-01 | 6.13E-01 | 8.90E-01 | 9.30E-01 | 0.16667 | -0.075 : 0.423 |
| rs17154155_ALT | -0.04608 | 4.96E-01 | 7.28E-01 | 8.85E-01 | 9.30E-01 | 0.08127 | -0.122 : 0.284 |

| | | | | | | | |
|---|---|---|---|---|---|---|---|
| rs1404312 | -0.04521 | 7.15E-01 | 9.03E-01 | 8.72E-01 | 9.30E-01 | -0.19578 | -0.454 : 0.064 |
| rs35660939 | -0.06731 | 5.31E-01 | 7.54E-01 | 9.58E-01 | 9.69E-01 | -0.09644 | -0.37 : 0.167 |
| rs11771152 | -0.00454 | 9.64E-01 | 9.89E-01 | 9.48E-01 | 9.69E-01 | -0.00632 | -0.267 : 0.256 |
| rs11770907 | -0.00199 | 9.83E-01 | 9.89E-01 | 9.37E-01 | 9.69E-01 | 0.03896 | -0.211 : 0.292 |
| rs9649529 | -0.01129 | 7.63E-01 | 9.14E-01 | 9.90E-01 | 9.90E-01 | -0.05748 | -0.237 : 0.128 |

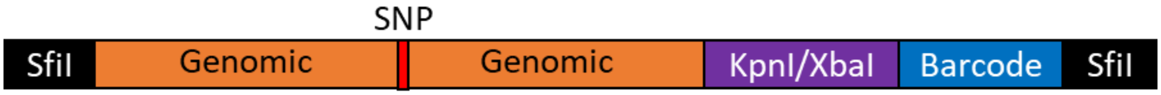


**Figure S1: MPRA Oligo Library Design.** Schematic of the oligonucleotide library used in the *CD36* MPRA. SfiI restriction sites generated by PCR were used to clone the oligo into the pMPRA1 backbone vector. *CD36* eQTL SNPs surrounded by 150bp of genomic context were located 5' to a KpnI/XbaI linker used to insert a minimal promoter-luciferase cassette derived from pNL3.2. 3' to the linker is a unique 40bp barcode used for identifying RNA regulated by the genomic seqeuence. Each allele has 40 barcodes associated with it.

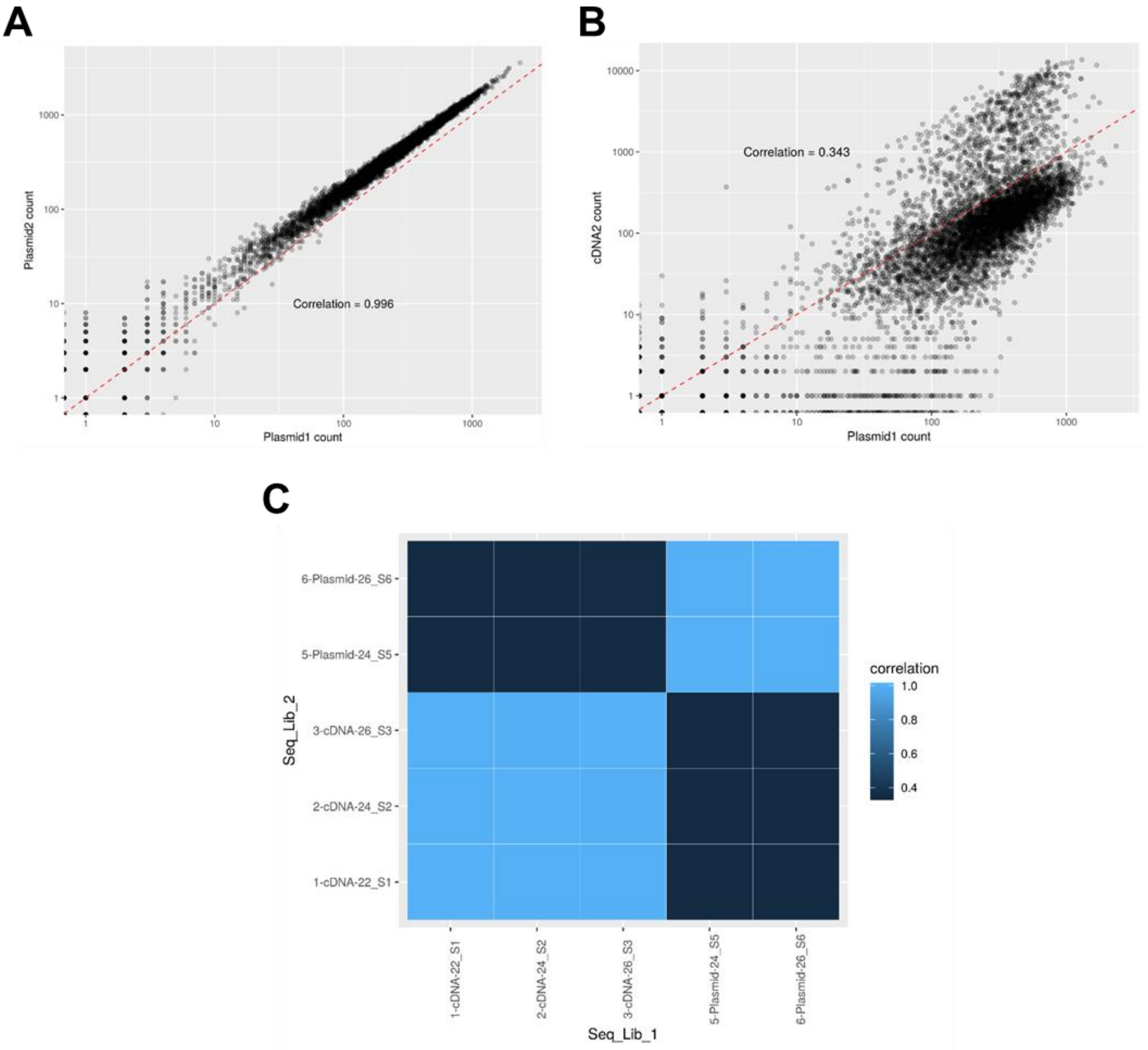


**Figure S2: Correlation of sequencing libraries:** (A) Scatterplot of barcode counts showing the correlation between the two sequencing samples prepared from the plasmid library. (B) A similar plot showing the expectedly lower correlation between one plasmid sample and a cDNA sample. Because the barcodes need to go through a transcription step to be present in the cDNA sample, these counts are inherently more noisy and less correlated with their plasmid barcode inputs. C) A heatmap showing the pairwise correlation values between all five samples in the experiment.

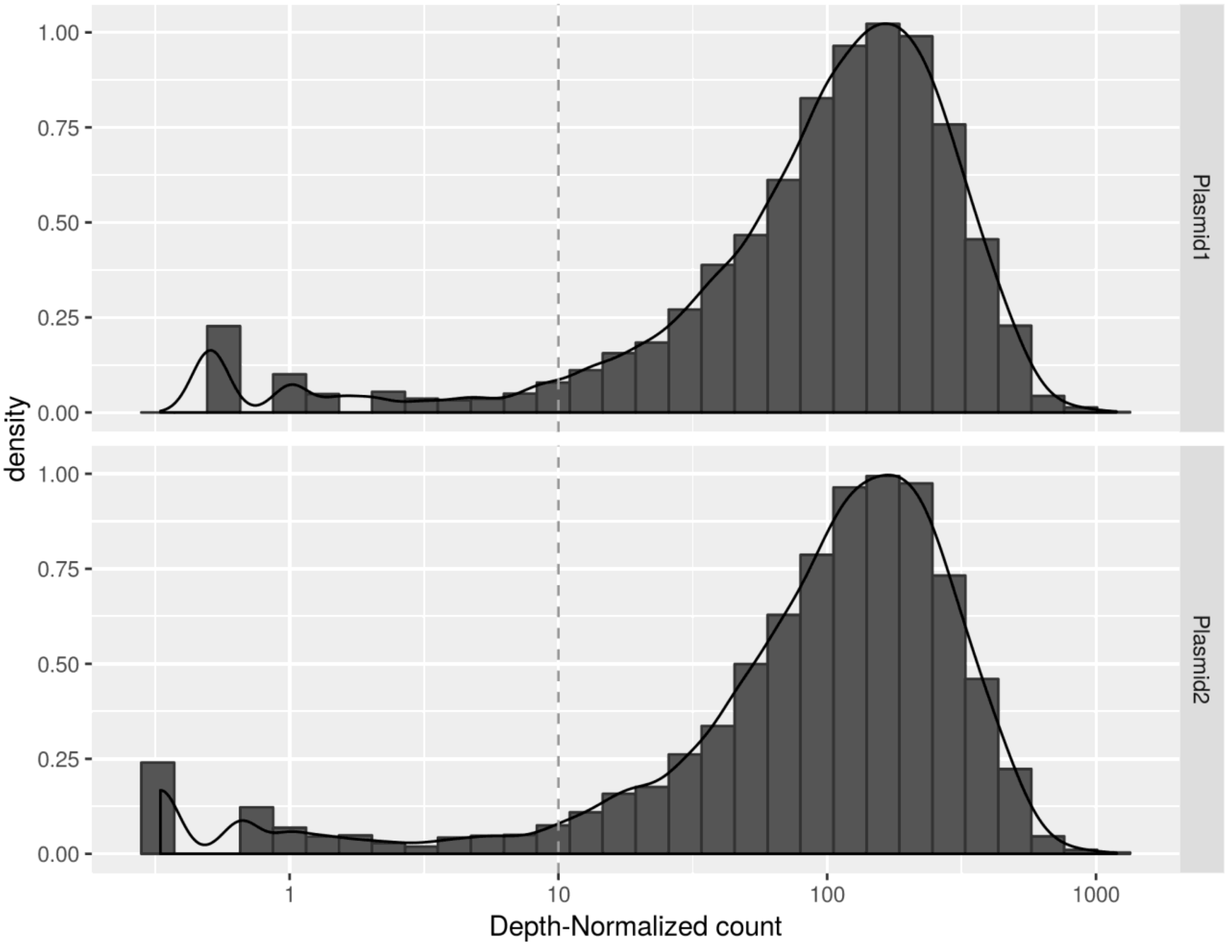


**Figure S3** – **Plasmid Library Barcode Representation.** A (A) histogram and (B) density function showing the abundance of DNA barcodes in the plasmid libraries. Barcodes present at very low levels in the plasmid library will cause the RNA output to be highly variable. A minimum plasmid library abundance is chosen by inspection to use as a cutoff. Barcodes that are not present above this level will be discarded from the downstream analysis. The counts are first normalized for sequencing depth by dividing through by the respective sample number of reads and multiplying by one million. A dashed line indicates the selected depth-normalized count cutoff.

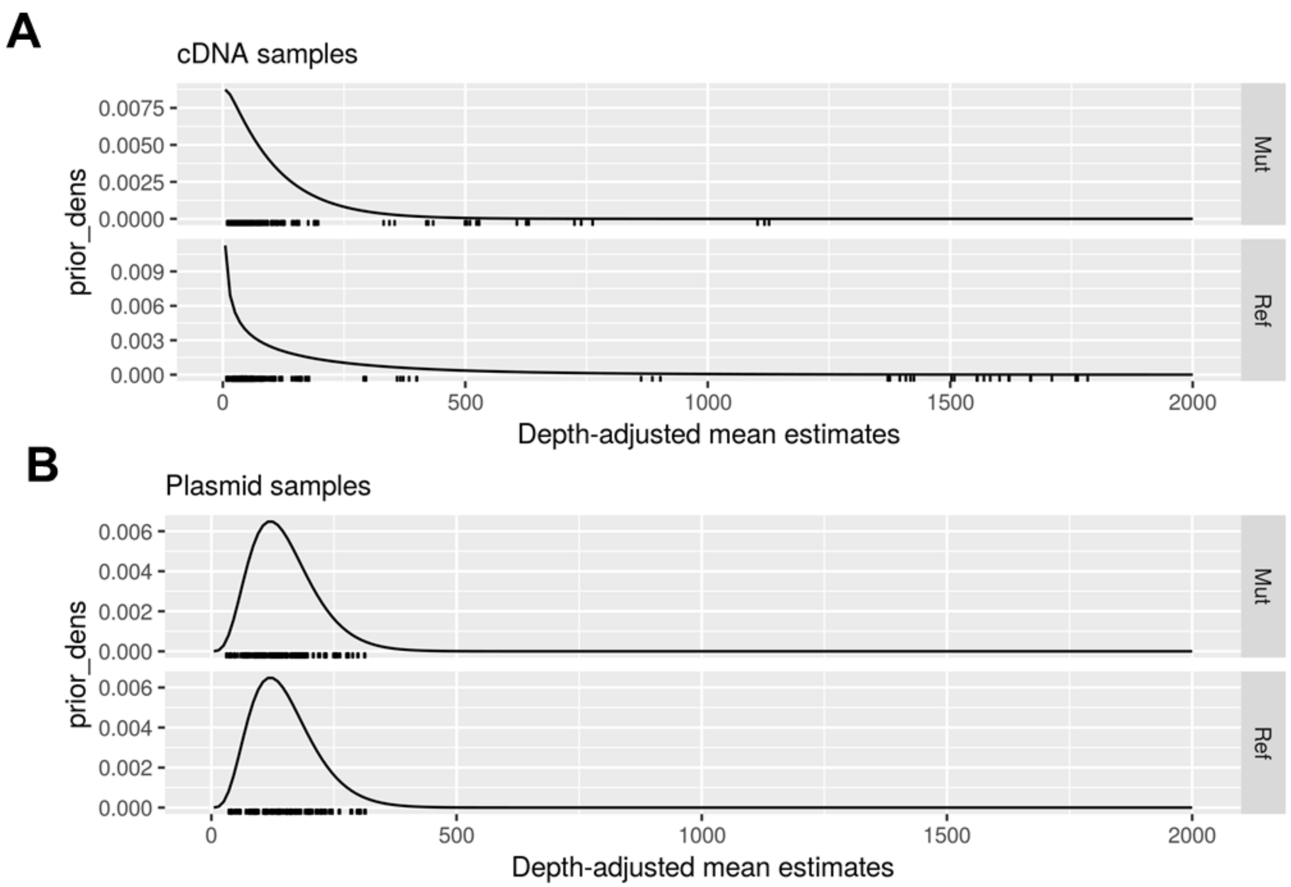


**Figure S4** – **Negative Binomial Mean Parameter Priors:** Density functions of the empirically estimated gamma priors for (A) the cDNA samples and (B) the plasmid library samples

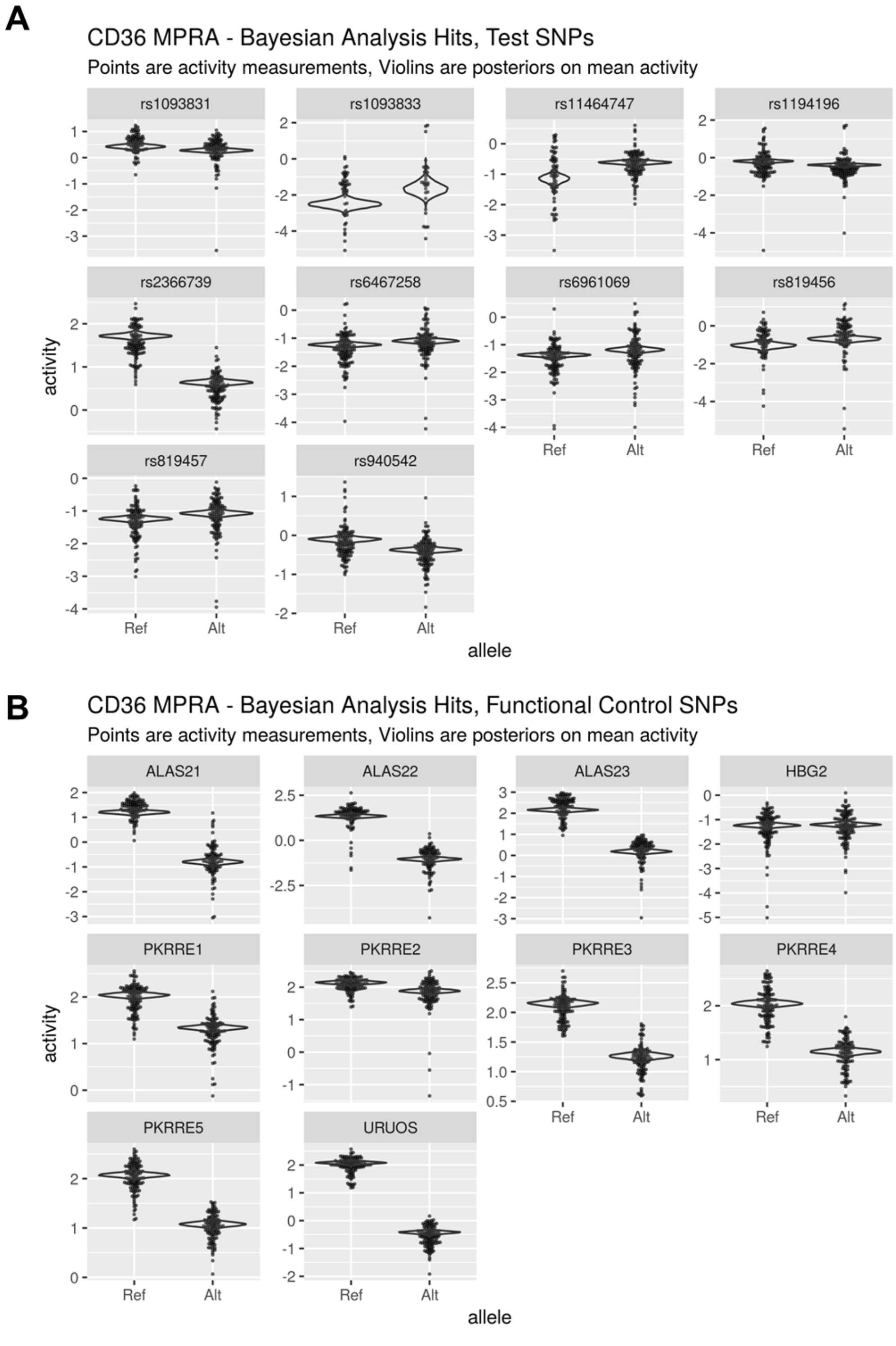


**Figure S5 – Combined activity scatterplots and posterior plots**: Points show activity measurements for individual barcodes, while violins show the posterior on the mean activity level for each allele from the Bayesian model for (A) the detected functional test SNPs and (B) the Controls.

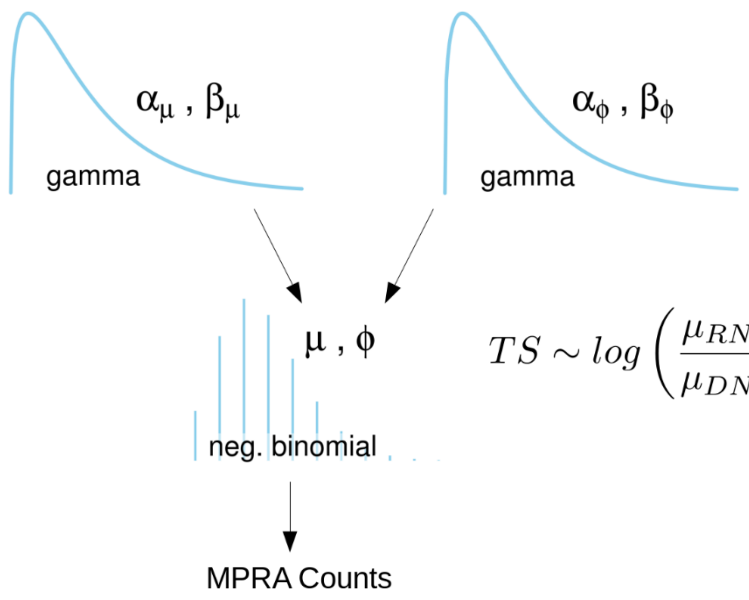


$$TS \sim log\left(\frac{\mu_{RNA,mut}}{\mu_{DNA,mut}}\right) - log\left(\frac{\mu_{RNA,ref}}{\mu_{DNA,ref}}\right)$$

**Figure S6** – Kruschke diagram showing the generative model underlying the Bayesian analysis. The mean parameters for the separate alleles and nucleic acids are aggregated after fitting the model into a posterior on transcription shift that is used to identify functional SNPs.
